# Ploidy variation and spontaneous haploid-diploid switching of *Candida glabrata* clinical isolates

**DOI:** 10.1101/2022.06.02.494626

**Authors:** Qiushi Zheng, Jing Liu, Juanxiu Qin, Bingjie Wang, Jian Bing, Han Du, Min Li, Fangyou Yu, Guanghua Huang

## Abstract

The human fungal pathogen *Candida glabrata* is phylogenetically closely related to *Saccharomyces cerevisiae*, a model eukaryotic organism. Unlike *S. cerevisiae* with both haploid and diploid forms and a complete sexual cycle, *C. glabrata* has long been considered a haploid and asexual species. In this study, we analyzed the ploidy states of 500 clinical isolates of *C. glabrata* from four Chinese hospitals and found that approximately 4% of the isolates were in or able to spontaneously switch to an aneuploidy (genomic DNA: 1N-2N), diploidy (2N), or hyperdiploid (>2N) form under *in vivo* or *in vitro* conditions. Stable diploid-form cells were identified in 3% of the isolates (15/500). Of particular interest, one clinical strain only existed in the diploid form. Multilocus sequence typing (MLST) assays revealed two major genetic clusters (A and B) of *C. glabrata* isolates. Most of the isolates (70%) from China belonged to the A cluster, whereas most of the isolates from other countries (such as Iran, Japan, USA, and European countries) belonged to the B cluster. Further investigation indicated that *C. glabrata* cells of different ploidy forms differed in a number of aspects, including morphologies, antifungal susceptibility, virulence, and global gene expression profiles. Additionally, *C. glabrata* could undergo spontaneous switching between the diploid and haploid form under both *in vitro* and *in vivo* conditions. Given the absence of an apparent sexual phase, one would expect that the ploidy shifts could function as an alternative strategy that promotes genetic diversity and benefits the ability of the fungus to rapidly adapt to the changing environment.

**Importance:** The human fungal pathogen *Candida glabrata* has long been thought to be a haploid organism. Here we report the population structure and ploidy states of 500 clinical isolates of *C. glabrata* from China. To our surprise, we found that the ploidy of a subset of clinical isolates varied dramatically. Some isolates were in or able to switch to an aneuploid, diploid, or hyperdiploid form. *C. glabrata* cells with different ploidy differed in a number of biological aspects, including morphologies, antifungal susceptibility, virulence, and global gene expression profiles. Given the absence of an apparent sexual phase in this fungus, we propose that ploidy switching could be a rapid adaption strategy to environmental changes and could function as an alternative strategy of sexual reproduction.

## 1. Introduction

Fungal infections became more and more common in clinical settings during the past several decades due to the increase in immunocompromised individuals (1, 2). *Candida glabrata*, a species phylogenetically closely related to *Saccharomyces cerevisiae*, is the second most important fungal pathogen behind *Candida albicans* in terms of infections (2, 3). *C. glabrata* differs from *C. albicans* in several aspects. First, the former often exists in the yeast form and is unable to form true hyphae or undergoes heritable phenotypic switching (such as white-opaque and white-gray-opaque transitions (4, 5)). Second, *C. glabrata* has been thought to contain a haploid genome, whereas *C. albicans* is an “obligate” diploid fungus. Third, *C. glabrata* often has a higher resistance to antifungal drugs, such as fluconazole. A previous study demonstrates that approximately 20% of *C. glabrata* isolates developed resistance to fluconazole during therapy (6). Fourth, a parasexual reproduction process has been reported in *C. albicans*. Despite similar mating type loci to those of *S. cerevisiae*, the process of sexual reproduction has not been observed in *C. glabrata*.

Genomic plasticity is an adaptive strategy to hostile environments and is often associated with fungal pathogenesis and antifungal resistance in many pathogenic fungi. It has been reported that the generation of aneuploidy is associated with the rapid evolution of new traits in *C. albicans, Cryptococcus neoformans*, and *Saccharomyces cerevisiae* (7-10). We recently found that the emerging fungal pathogen *Candida auris* could also undergo ploidy alterations to adapt to environmental changes (11). Changes in ploidy state, including the generation of aneuploidy, have a profound effect on many physiological aspects, such as cell size, growth rate, genomic stability, and transcriptional output (7-10). The number and size of chromosomes vary dramatically, and rearrangements of the genome occur frequently in clinical isolates of *C. glabrata* (12). These genomic changes have been found to be associated with both virulence and antifungal resistance of the fungus (13, 14). Genomic evidence of recombination indicating that *C. glabrata* could be able to undergo sexual reproduction or have a cryptic sexual cycle (15, 16). Given the lack of an observed/reported sexual cycle in *C. glabrata*, this genomic plasticity would be fundamentally important to its adaptation to the ever-changing host environment.

In this study, we sought to analyze the phenotypic and genetic diversity of clinical isolates of *C. glabrata*. By analyzing 500 clinical strains of *C. glabrata* isolated from four different Chinese hospitals, we discovered two major genetic clades (A and B). Most isolates from China belonged to clade A. To our surprise, approximately 4% of the isolates existed in or were able to spontaneously switch to an aneuploid (genomic DNA: 1N-2N), diploid (2N), or hyperdiploid (>2N) form. Of them, fifteen independent strains could exist in the stable diploid form. To our knowledge, this is the first report of the diploid form of clinical *C. glabrata* strains. We observed that cells with different ploidy forms exhibited different susceptibilities to antifungals, appearances on CuSO_4_-containing medium, and fungal burdens during infection. However, the direct relationship between these phenotypes and ploidy variations and the underlying genetic basis needs further investigation.

## 2. Materials and methods

### 2.1 Strains and culture conditions

Clinical isolates of *C. glabrata* were collected from several Chinese hospitals in Shanghai, Shenyang, and Guiyang in China. The detailed information on these isolates (original tissue sources, ploidy, hospital information, and information on associated patients) are presented in supplementary **Dataset S1**. All samples were plated on CHROMagar medium for initial identification of *Candida* species. The potential *C. glabrata* isolates were further verified by sequencing the ITS (internal transcribed spacer) region (primers used for PCR: ITS1: GTCGTAACAAGGTTTCCGTAGGTG, NL4: GGTCCGTGTTTCAAGACGG). YPD medium was used for regular growth of fungal cells. Phloxine B (5 μg/mL), which could stain certain types of colonies red or pink, was added to YPD medium. YPD + CuSO_4_ medium (1mM CuSO_4_) was used for phenotypic analysis of haploid, diploid, or hyperdiploid cells of *C. glabrata*.

### 2.2 Fluorescence-activated cell sorting (FACS) analysis

The content of genomic DNA of different isolates were analyzed using FACS analysis according to our previous description (17). *C. glabrata* cells were initially grown on YPD medium for 72 hours at 30°C. The cells were then inoculated and incubated in liquid YPD medium with shaking for overnight growth (at 30°C, 220 rpm). They were then reinoculated into fresh medium (initial OD_600_ = 0.3), cultured to logarithmic phase (OD_600_ = 1.8), and harvested for flow cytometry analysis. Briefly, fungal cells were washed with ddH_2_O, resuspended in 300 μL 1 x TE buffer (1.2114 g/L Tris, 0.29224 g/L EDTA, adjusted the pH to 8.0) and mixed with 700 μL 100% ethanol in 1.5 mL RNase-free tubes. The samples were placed on vortex mixer (Vortex-Genie 2) for at least two hours at ambient temperature. The fungal cells were then washed and resuspended with 1 x TE buffer and treated with RNase A (concentration: 0.3 mg/mL) overnight at 37°C. The cell samples were further treated with proteinase K (concentration: 0.03 mg/mL) and then incubated for two hours at 50°C. Fungal cells were collected, washed twice with 1 x TE buffer, and suspended in 1 x PBS. Cells were stained with propidium iodide (concentration: 25 μg/mL) and used for analysis of genomic DNA content. The amount of at least 30,000 cells of each sample were detected on a FACS Caliber. The software FlowJo 10.4 was used for analyzing the data.

### 2.3 Multi-locus sequence typing (MLST) analysis

Six gene loci were used for MLST analysis. The primers used: *FKS* (5’-GTCAAATGCCACAACAACAACCT, 3’-AGCACTTCAGCAGCGTCTTCAG), *LEU2* (5’-TTTCTTGTATCCTCCCATTGTTCA, 3’-ATAGGTAAAGGTGGGTTGTGTTGC), *NMT1* (5’-GCCGGTGTGGTGTTGCCTGCTC, 3’-CGTTACTGCGGTGCTCGGTGTCG), TRP1 (5’-AGCACACAGGGATTGTTGTA, 3’-GACCAGTCCAGCTTTTCAC), *UGP1*(5’-TTTCAACACCGACAAGGACACAGA, 3’-TCGGACTTCACTAGCAGCAAATCA), and *URA3* (5’-AGCGAATTGTTGAAGTTGGTTGA, 3’-AATTCGGTTGTAAGATGATGTTGC). The primers were designed based on previous studies (18, 19). We sequenced the six genes of 70 *C. glabrata* strains (including 20 isolates with ploidy variation and 50 haploid). The 50 haploid strains were randomly selected from the four main sources listed in **Table 1** (26 from the respiratory tract, 10 from urine and feces, 8 from body fluid, and 6 from the genital tract). The sequences of 124 *C. glabrata* strains from Iran and USA were retrieved from the NCBI GeneBank database. Sequences were aligned using mafft v7.015b (20). Six alignments were concatenated into a single alignment. The maximum likelihood (ML) phylogenetic tree was generated using the program RAxML v7.3.2 (21) and the GTRGAMMA model with 1000 bootstraps.

**Table 1.** Percentages of clinical samples of *C. glabrata* containing varied ploid forms.

| Source | Number of samples | Percentage (%) of samples containing |  |  |  |
| --- | --- | --- | --- | --- | --- |
|  |  | Haploid | Diploid | Aneuploid or hyperdiploid | Diploid or aneuploid or hyperdiploid |
| Respiratory tract | 272 | 99.63% (271) | 4.04% (11) | 1.84% (5) | 4.41% (12) |
| Genital tract | 39 | 100.00% (39) | 0% (0) | 5.13% (2) | 5.13% (2) |
| Body fluid | 45 | 100.00% (45) | 2.22% (1) | 6.67% (3) | 6.67% (3) |
| Urine and feces | 144 | 100.00% (144) | 2.08% (3) | 0.69 % (1) | 2.08% (3) |
| **All** | **500** | **99.80% (499)** | **3.00% (15)** | **2.20% (11)** | **4.00% (20)** |

**Notes:** Respiratory tract (samples from sputum, bronchoalveolar lavage fluid, and oral swabs were included); body fluid (the abdominal fluid, ascites, blood, pleural fluid, and pus were included; urine and feces (urine, feces, associated catheters, and drainages were included). The percentages represent the ratios of samples containing certain types of ploid cells. The numbers in brackets represents the numbers of samples containing certain types of ploid cells.

### 2.4 Minimum inhibitory concentration (MIC) assay

MICs of *C. glabrata* strains were determined according to the NCCLS document M27-A2 and previous investigation (22). Fungal cells were initially grown on YPD medium at 30°C for two days and then collected and washed with ddH_2_O. Approximately 500 cells were incubated in 200 μL liquid RPMI-1640 medium (w/v, 1.04% RPMI-1640, 3.45% MOPs, adjusted the pH to 7.0) using a 96-well U-bottom microplate. A series concentrations of itraconazole (0.0625, 0.125, 0.25, 0.5, 1, 2, 4, and 8 μg/mL) was tested. Three biological repeats were performed for each isolate. *Candida parapsilosis* ATCC22019 and *Pichia kudriavzevii* ATCC6258 served as quality controls. The microplates were incubated at 35°C for 24 hours. The growth status of cells at different concentrations was determined using a microplate reader. The MIC value was determined when the growth state and turbidity of the tested wells was more than 50% lower than that of the control.

### 2.5 Animal experiments

All animal experiments were approved by the Animal Care and Use Committee of Fudan University. Four mice (four-week-old female BALB/c) were used for systemic infection for each strain of *C. glabrata*. Four mice were housed in a cage with an adequate supply of food and water. All the mice were stayed in the animal facility for a week of acclimation before used for infection. *C. glabrata* strain used: RJ155 (haploid control), FK83-1 (haploid), and FK83-2 (diploid). FK83-1 and FK83-2 are isogenic isolates. The haploid-only isolate (RJ155) served as the control. Fungal cells were initially grown on YPD medium at 30°C for two days and then collected, washed, counted, and diluted with 1 x PBS. Each mouse was injected with 2 × 10^7^ fungal cells in 250 μL 1 x PBS via the lateral tail vein. After 24 hours of infection, the mice were humanely killed. The brain, liver, spleen, lung, and kidney tissues were collected, weighed, ground up, and plated on YPD medium for the fungal burden and morphological assays. The YPD medium was supplemented with ampicillin sodium (final concentration 100 μg/mL) and kanamycin sulfate (final concentration 50 μg/mL).

### 2.6 RNA-seq analysis

Haploid and diploid cells of *C. glabrata* (FK83) were incubated in liquid YPD medium for overnight growth at 30°C and re-inoculated into fresh liquid YPD medium with the initial OD_600_ of 0.3. Fungal cells were grown to the logarithmic growth stage and collected for total RNA extraction. Library preparation and sequencing were by the company Berry Genomics (Beijing, China). Briefly, approximately 3 G bases of reads were obtained by sequencing each library. Three biological repeats were performed for each strain. The sequence data of *C. glabrata* CBS138 (GenBank accession: GCA_000002545.2) were used as the reference. Clean reads were mapped to the reference sequence using the software HiSat2 v2.0.5 with default parameters. Transcriptional expression levels of different samples were estimated with StringTie v1.3.3b with default parameters (23). Differentially expressed genes were analyzed with the DESeq2 R package (24). Differential expressed genes must satisfy two criteria: (1) fold change ≥2; (2) false discovery rates (FDRs) ≤0.05.

### 2.7 Microscopy assays

The colony phenotypes were observed by using Stereomicroscope (NSZ-810+Dig1600, Shanghai Wentek Photonics Technology Co., Ltd, Shanghai, China) and the cellular morphologies were observed using Leica DM2500 (Wetzlar, Germany). To measure the cell parameters, fungal cells were photographed under 40 x objective lens, the cell size was measured (20 cells/strain).

### 2.8 Statistical methods

Student’s *t*-test (two tailed) and F-test were used for Figure 6. F-test was used to compare and judge the homogeneity of variance. Equal variance was considered if the result > 0.05. Cohen’s *d* was used to measure the standardized effect size for *t*-tests. 0.2 ≤ d < 0.5 indicates a small effect; 0.5 ≤ d < 0.8 indicates a medium effect; and d ≥ 0.8, indicates a large effect.

### 2.9 Spontaneous ploidy switching assays

For *in vitro* ploidy switching assay, haploid cells of two strains (FK83 and FK181) were initially grown on YPD medium at 30°C for two days. Fungal cells of homogeneous colonies (white) were replated on YPD + phloxine B medium at 30°C for seven days. Approximately 100 cells were plated on each YPD plate. The cells of colonies appeared pink or red were replated on YPD + phloxine B medium and cultured for seven days at 30°C. Flow cytometry analysis was performed to verify the ploidy state. The frequency of white-to-pink/red colony switching was regarded as the ploidy switching frequency.

For *in vivo* ploidy switching assay, haploid cells of two strains (FK83 and FK181) were initially grown on YPD medium at 30°C for two days. Fungal cells were collected, washed, and diluted with 1 × PBS. Cells were used for animal experiments as described in Section 2.5. Fungal cells were recovered from five organs and plated on YPD medium with ampicillin sodium (final concentration 100 μg/mL) and kanamycin sulfate (final concentration 50 μg/mL). The colonies with alternative coloration (gray) were replated and cultured on YPD + phloxine B medium at 30°C for seven days. Flow cytometry analysis was performed to verify the ploidy state. The frequency of ploidy switching was determined.

## 3. Results

### 3.1 Collection of clinical isolates of *C. glabrata*

We collected 500 clinical isolates of *C. glabrata* from four different hospitals in China. Of them, 211 strains were collected from Renji Hospital (Shanghai), 98 from Shanghai Pulmonary Hospital (Shanghai), 150 from the First Hospital of China Medical University (Shenyang), and 41 from Guiyang Hospitals (Guiyang). The clinical information for the associated isolates is summarized in supplementary **Dataset S1**. CHROMagar medium was used for initial identification of *Candida* species. *C. glabrata* isolates were verified by sequencing the ITS region. All associated patients were adults (from 18 to 98 years old) including 295 male (59.0%) and 205 female (41.0%) patients. Among the *C. glabrata* strains, 281 (56.2%) were isolated from patients more than 70 years old, 154 (30.8%) from those 50 to 70 years old, and 65 (13.0%) from those less than 50 years old. The original sources of *C. glabrata* isolates included sputum (n=263, 52.6%), urine (n=92, 18.4%), genital secretions (n=39, 7.8%), blood (n=37, 7.4%), feces (28, 5.6%), and others (n=39, 7.8%). The strains represented all isolates of a certain period from the corresponding hospital (**Dataset S1**).

### 3.2 Discovery of the diploid form of *C. glabrata*

*C. glabrata* cells normally formed purple or light purple colonies on CHROMagar medium (**Figure S1**). Some colonies, which were formed by plating the clinical specimen and identified as *C. glabrata*, exhibited much darker coloration than the rest. The cells of these colonies were notably larger than normal *C. glabrata* cells (**Figure S1**). We then patched these colonies on YPD medium and examined the genomic DNA content using Flow cytometry analysis (FACS) assays. As shown in **Figure S1**, *C. glabrata* formed distinct colonies with different levels of coloration. The cells of the darker colonies had a diploid genome, whereas the cells of the regular colonies had a haploid genome. Our findings imply that natural isolates of *C. glabrata* can exist in the diploid form in the human host.

### 3.3 Ploidy variation of *C. glabrata* isolates

We next asked whether ploidy variation was a general feature of clinical isolates. To facilitate the identification of different ploidy forms of *C. glabrata*, we developed a method by culturing fungal cells on YPD agar containing phloxine B (5 μg/mL) for six days at 30°C. As shown in **Figure 1**, colonies formed by diploid, aneuploid, or hyperdiploid cells often exhibited a pink or red coloration, whereas colonies formed by haploid cells were shiny and white. Strain JX1092 was a typical haploid strain, the cells of which were relatively small and the colonies white in the presence of phloxine B (**Figure 1A**).

**Figure 1.**
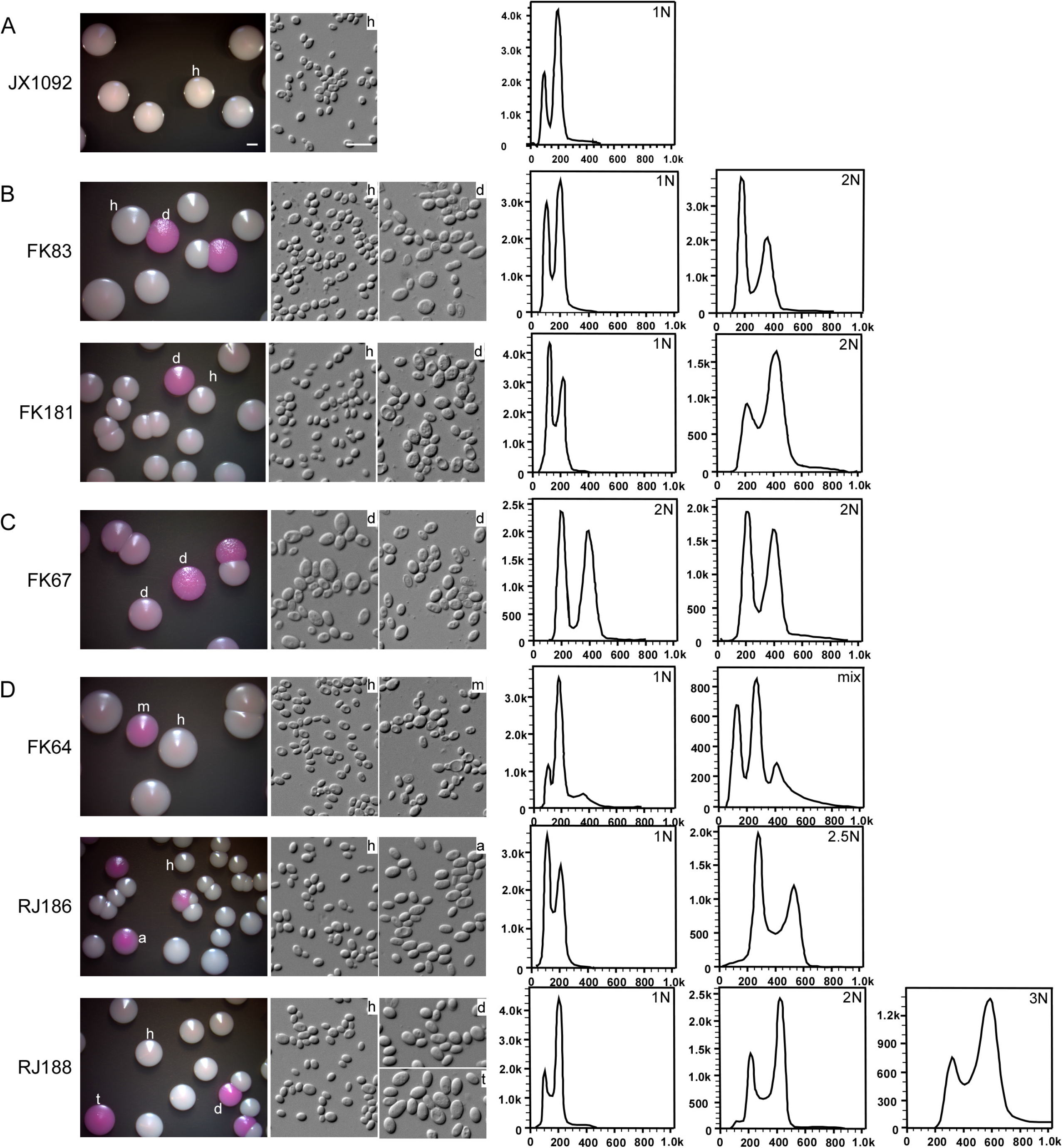
Colony and cellular morphologies of representative isolates of *C. glabrata* containing haploid, diploid, aneuploid, or hyperdiploid cells. Cells of *C. glabrata* isolates were initially grown on YPD medium for three days. Cells of a representative colony were replated on YPD medium (containing 5 μg/mL of phloxine B) and incubated at 30°C for six days. FACS analyses of colonies with different appearances were performed to determine the genomic DNA content. **(A)** JX1092, a stable haploid isolate; **(B)** FK83 and FK181, two isolates containing both haploid and diploid cells; **(C)** FK67, a stable diploid isolate; **(D)** FK64, RJ186, and RJ188, three isolates containing haploid, aneuploid or hyperdiploid cells. h, haploid; d, diploid; m, mix; a, aneuploid; t, triploid. Scale bar for colonies, 2 mm; Scale bar for cells, 10 μm.

Using this method, we analyzed the collected 500 clinical isolates of *C. glabrata*. Most isolates only formed regular white colonies on YPD medium in the presence of phloxine B. Cells of the colonies were also relatively small. We found that 20 isolates (4.0%) had two or more phenotypes, including white, light pink, and red colonies (**Figure 1B** and **1D**). All of the colonies with different phenotypes were verified and identified as *C. glabrata* by sequencing the ITS region. Microscopic examination demonstrated that the cell size (diameter) of light pink or red colonies (7.5 ± 0.9 μm) was significantly larger than that of white colonies (3.1 ± 0.6 μm). Approximately 20 representative cells were examined for each isolate. FACS analysis confirmed that fungal cells of the 20 isolates with different colorations had distinct genomic DNA content (including diploidy, aneuploidy, and hyperdiploid). The colony and cellular morphologies and FACS results of several examples with different ploidy forms are presented in **Figure 1**. A summary of the ploidy forms of the 500 isolates is presented in **Table 1** and supplementary **Dataset S1**. Interestingly, although strain FK67 had two different colony phenotypes, cells of both colony types were diploid (**Fig 1C**). This isolate could only exist as the diploid form during infection. However, the coloration levels of both morphologies (pink or red) were higher than that of the haploid form (white). Strain FK67 could represent a special case and reflect the diversity of morphology of *C. glabrata*. Therefore, a combined assay of the test of genomic content and morphological observation should be performed to accurately determine the ploidy state. Of the 20 samples containing cells with altered ploidy forms (non-haploid), 15 had diploid, 11 had aneuploidy/hyperdiploid, and 6 had both diploid cells and aneuploid/hyperdiploid cells. All isolates contained cells of the haploid form, except for strain FK67, which was isolated from the respiratory tract and only existed as the diploid form. The ratio of isolates with altered ploidy forms from the respiratory tract, genital tract, body fluid and tissue, urine, and feces was 4.4%, 5.1%, 6.7%, and 2.1%, respectively. The hypergeometric distribution demonstrated that the samples from the respiratory tract had a weakly higher ratio of the diploid form than the other sources (P=0.07). Taken together, although the ratio was different perhaps due to the small size of samples, isolates with at least one altered ploidy (>1N) could be found in all four tissue sources (**Table 1**).

### 3.4 Genotyping of clinical isolates of *C. glabrata*

We next performed a genotyping analysis using the multi-locus sequence typing (MLST) assay. The 20 isolates containing diploid or aneuploid/hyperdiploid cells and 50 haploid-only isolates (randomly selected from the four tissue sources) were examined. Six MLST loci (*URA3, UGP1, TRP1, NMT1, LEU2*, and *FKS*) were genotyped according to previous reports (18, 19). As shown in **Figure 2**, the 70 isolates were clustered into two major genetic clades (A and B). Most strains belonged to clade A (70%, 49/70), which included 17 (85%, 17/20) samples with an altered ploidy (highlighted in blue, **Figure 2**). Overall, the percentage of strains with changed ploidy was two-fold greater in clade A (34.7%, 17/49) than clade B (14.3%, 3/21), implying a bias distribution of samples with an altered ploidy form between the two genetic clades. However, this genetic distribution was not significantly related to the tissue sources and certain hospitals.

**Figure 2.**
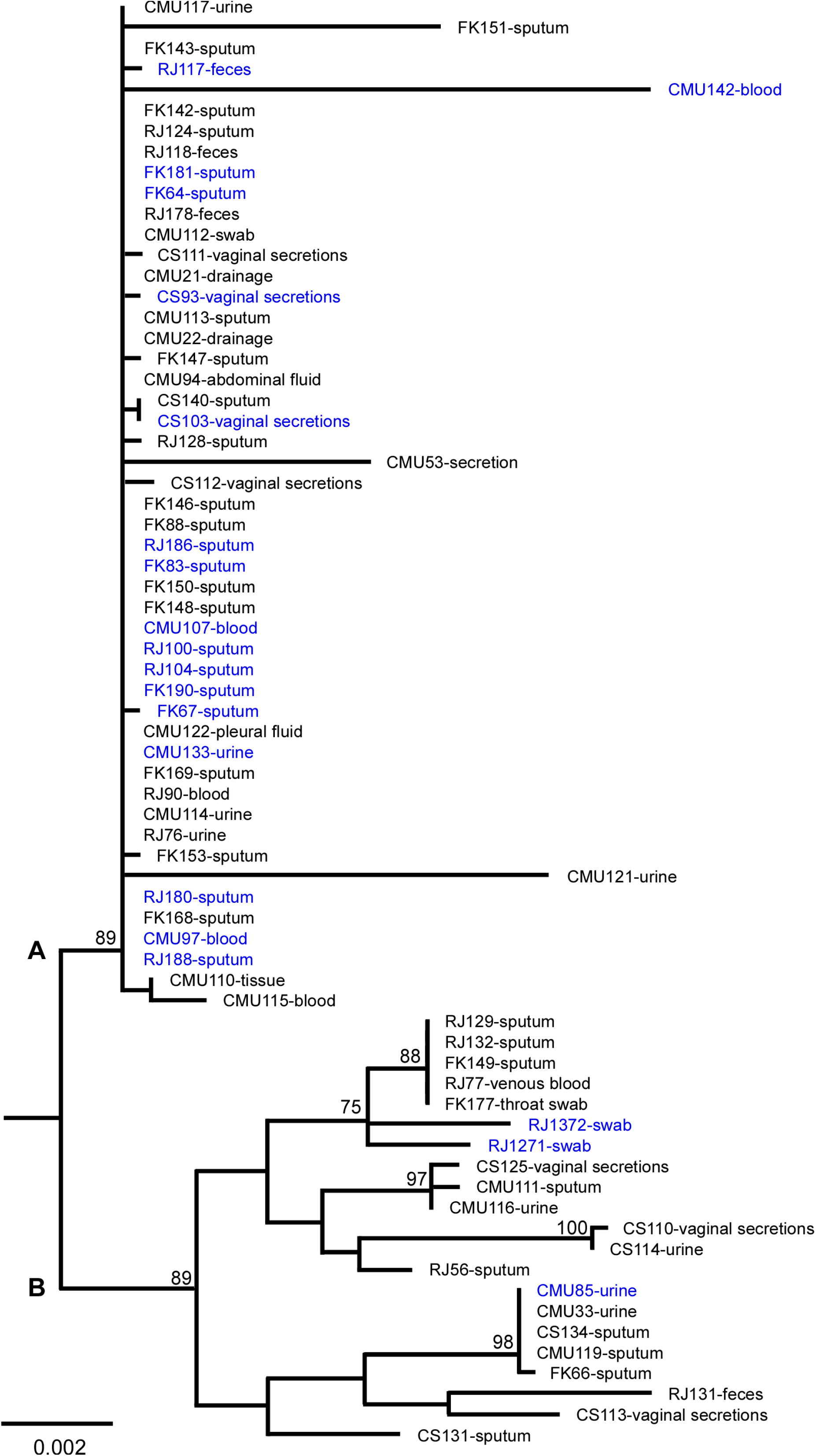
The population structure of 70 *C. glabrata* isolates from four Chinese hospitals (including 20 with variable ploidy and 50 haploid-only isolates). Multi-locus sequence typing (MLST) analysis was performed based on six genes (*FKS, LEU2, NMT1, TRP1, UGP1*, and *URA3*). The phylogenetic tree was constructed using the maximum-likelihood (ML) method. The General Time Reversible (GTR) model, Gamma distribution, and 1000 bootstraps were adopted. CMU: The first hospital of China medical university;RJ: Shanghai Renji hospital;FK: Shanghai pulmonary hospital;CS: Clinical Strains (Guiyang Hospitals). According to MLST analysis, *C. glabrata* strains were divided into two clusters (A and B). The strain names and sources of isolation were included and presented in the tree. The names in blue indicate the clinical strains with diploidy and/or hyperdiploidy forms. Horizontal bars represent expected number of substitutions per site.

To examine whether there was a difference in genetic distributions between *C. glabrata* isolates from China and other countries, we retrieved the reported sequences of the six MLST loci of 124 non-Chinese strains from GenBank (www.ncbi.nlm.nih.gov/genbank, accession numbers: KX187005-KX187304; AY771006-AY771209, and KT763084-KT763323; strains from Iran, Japan, USA, some South American and European countries) [16-18]. The population structure analysis demonstrated that most non-Chinese strains were clustered into clade B (black, 108/124), whereas most Chinese isolates belonged to clade A (**Figure S2**).

### 3.5 Spontaneous ploidy switching in *C. glabrata in vitro* and *in vivo*

Since many samples from the same tissue of a patient could exist in two or more types of ploidy forms, we suspected that *C. glabrata* could undergo spontaneous switching between different ploidy forms under *in vitro* or *in vivo* conditions. To test this hypothesis, we performed switching assays on YPD medium containing phloxine B. Haploid cells of two strains (FK83 and FK181) were initially plated on the medium. After seven days of growth, approximately 0.3%-0.6% colonies appeared pink or red. The pink or red colonies were then replated on YPD medium. FACS analysis demonstrated that these colonies were an intermediate form containing haploid and diploid mixed cells. After another round of plating, homogeneous diploid colonies formed (**Figure 3**). To test whether diploid cells of *C. glabrata* could return to the haploid form, we cultured diploid cells of strains FK83 and FK181 on the same medium plates. Interestingly, diploid cells first switched back to the intermediate form (with a frequency of 1.0%-1.4%) and then to the haploid form (**Figure 3**). These findings demonstrate that only a portion of cells initially switched to the alternative form in the colonies.

**Figure 3.**
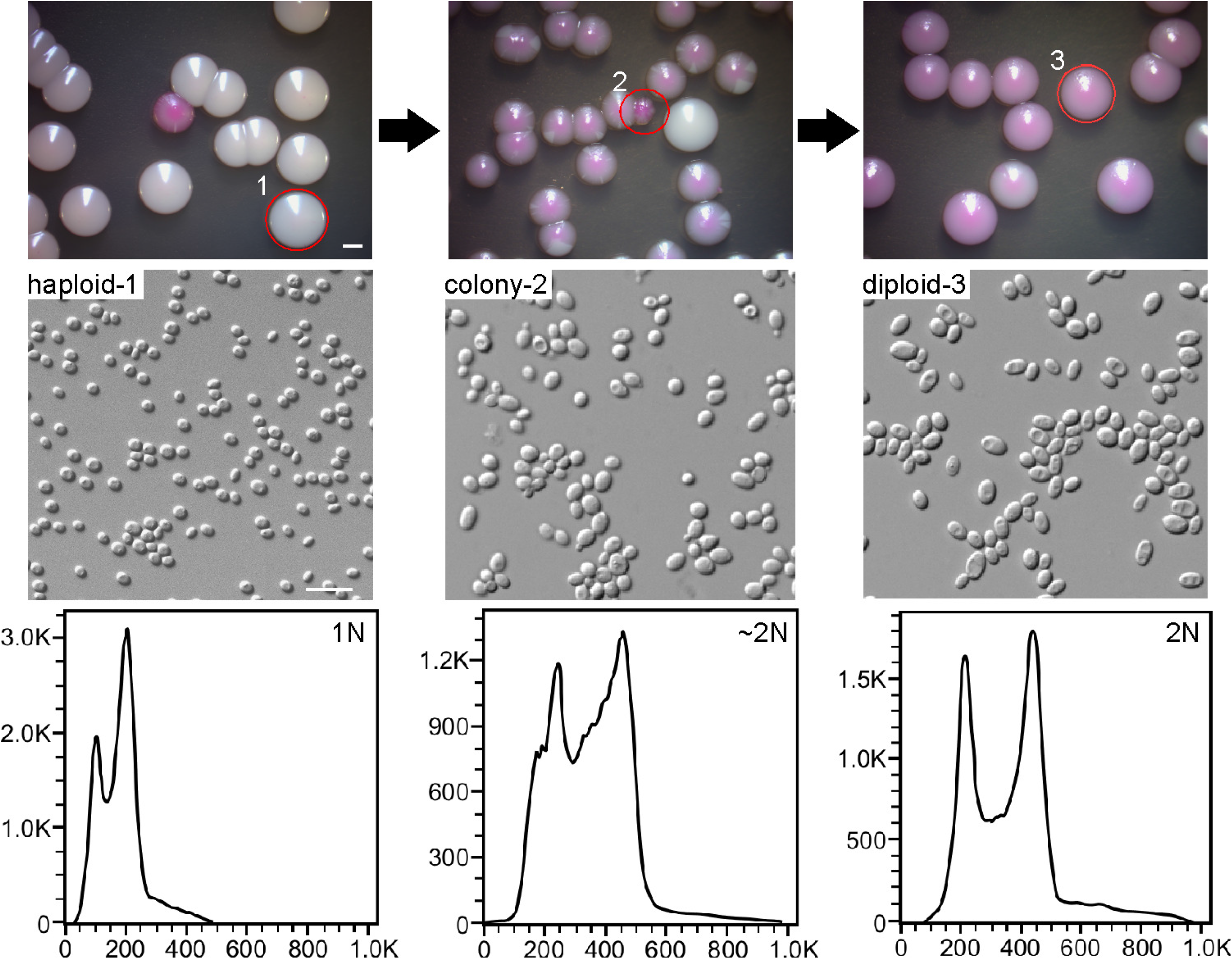
Haploid-diploid switching of *C. glabrata* on YPD medium. Strain FK83 was used. Haploid cells were initially grown on YPD medium for three days. Cells of a homogeneous colony (white) were plated on YPD medium (containing 5 μg/mL of phloxine B) and incubated at 30°C for six days. Pink or red colonies were replated on YPD medium, incubated at 30°C for six days, and subject to FACS analysis. When haploid cells were plated, colonies of an intermediate form developed that contained a mixture of both haploid and diploid cells. The switching frequency from haploid to mixed/diploid form was ∼0.6%, whereas the switching frequency from diploid to haploid was ∼1.0%. Scale bar for colonies, 2 mm; scale bar for cells, 10 μm.

To mimic the situation of human infection, we next performed ploidy switching assays using a mouse infection model. The mice were systemically infected with haploid or diploid cells of *C. glabrata* strains FK83 and FK181 through the vein tail injection. After 24 hours of infection, the mice were humanely killed, and fungal cells were recovered from the liver tissues and then plated on YPD medium without phloxine B. Some gray colonies (0.1%-0.3%) were observed and replated on YPD medium containing phloxine B (**Figure 4**). After seven days of growth at 30°C, both diploid (red) and haploid (haploid) colonies formed in both strains. However, we did not observe diploid-to-haploid conversion in either strain in the systemic mouse infection model (frequency <0.2%).

**Figure 4.**
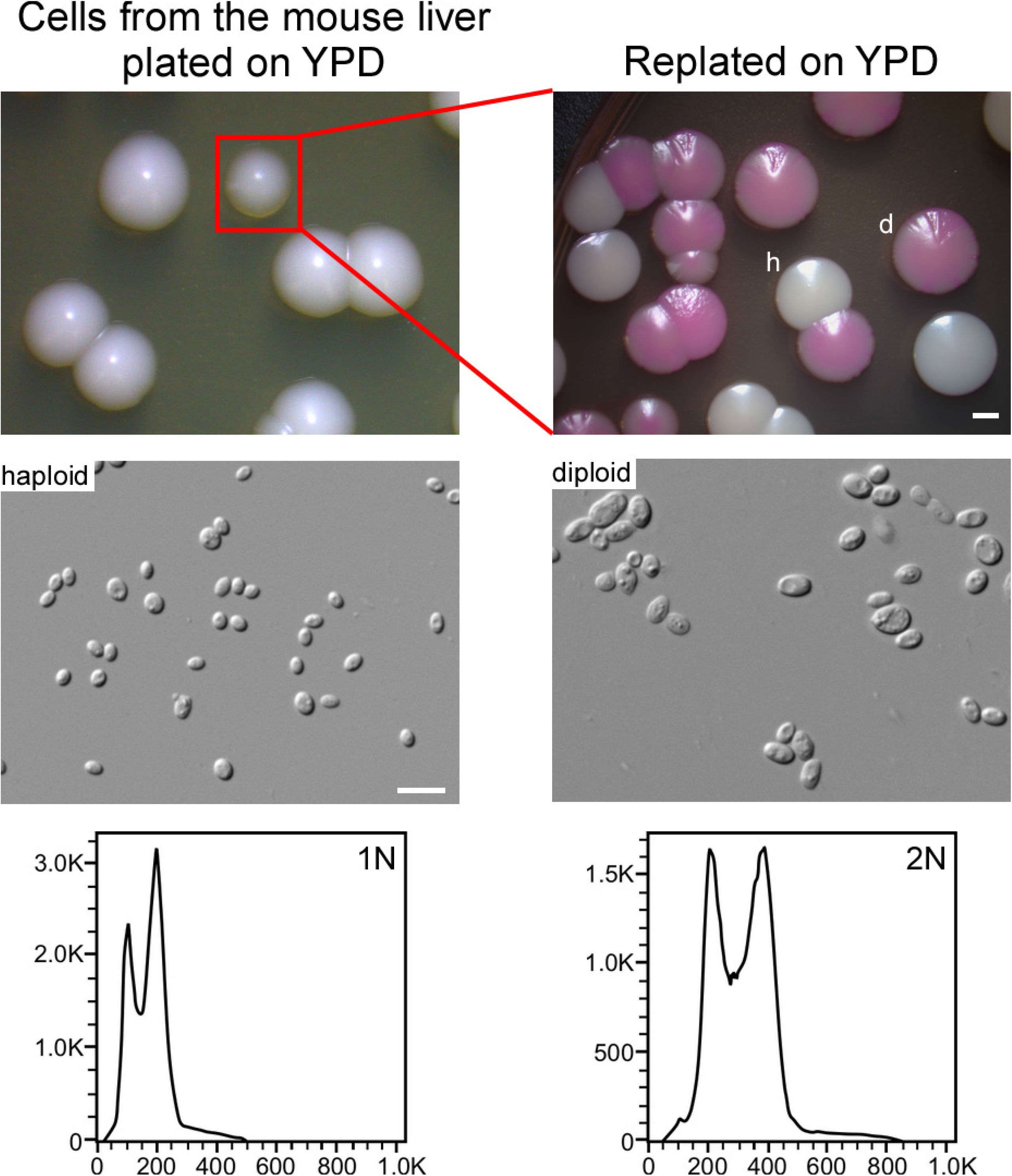
Haploid-diploid switching of *C. glabrata* in a mouse infection model. Strain FK83 was used. Haploid cells were injected into the mice via the tail vein. Fungal cells were recovered from the liver 24 hours post-infection, replated on YPD medium, and incubated at 30°C for three days (left panel). Colonies with darker coloration were replated on YPD medium (containing 5 μg/mL of phloxine B, right panel) and incubated at 30°C for six days. FACS analysis was performed to determine the genomic DNA content. The switching frequency from haploid to diploid form was ∼0.3%, whereas the switching frequency from diploid to haploid was <0.2%. Scale bar for colonies, 2 mm; scale bar for cells, 10 μm.

Taken together, we found that *C. glabrata* is able to undergo ploidy changes under both *in vitro* and *in vivo* conditions, although the switching frequencies are relatively low.

### 3.6 Cells of different ploidy forms show distinct phenotypes on CuSO_4_-containing medium

When cultured on YPD + CuSO_4_ medium, haploid *C. glabrata* strains could form distinct colonies with different colorations due to the distinct ability to convert CuSO_4_ into CuS (25, 26). This ability has been reported to be associated with the virulence of *C. glabrata* (27). Five representative strains with at least two ploidy forms were spotted on YPD + CuSO_4_ and YPD + phloxine B medium plates and incubated at 30°C for seven days. As shown in **Figure 5**, the order of coloration was hyperdiploid > diploid > haploid on YPD + phloxine B medium, whereas it was the reverse on YPD + CuSO_4_ medium (haploid > diploid > hyperdiploid). These results imply that *C. glabrata* cells with distinct ploidy forms can differ in the cellular oxidation–reduction (redox) state.

**Figure 5.**
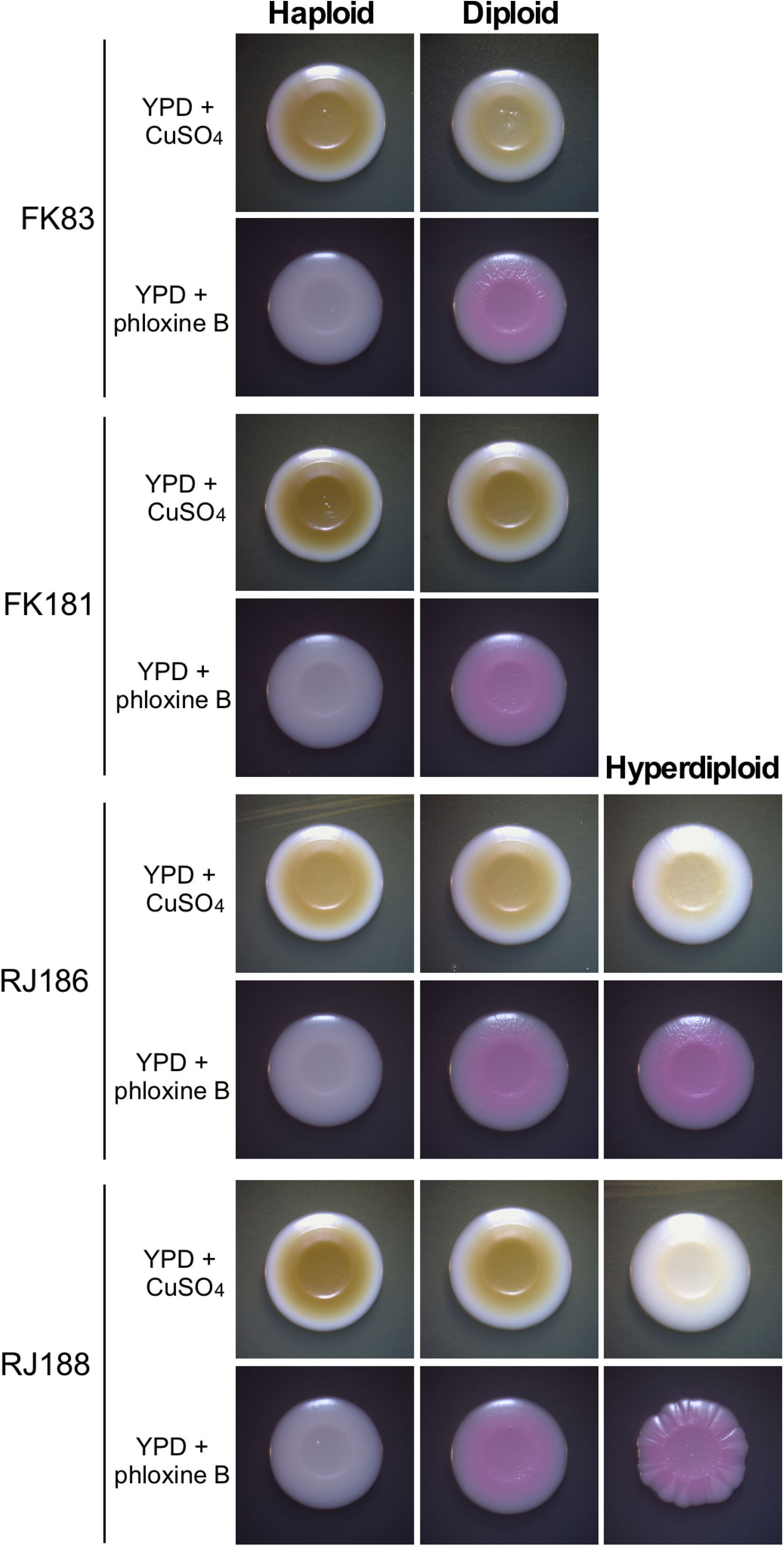
Phenotypes of haploid, diploid, and hyperdiploid cells of *C. glabrata* grown on YPD + CuSO_4_ and YPD + phloxine B media. *C. glabrata* cells were initially grown on YPD medium for three days. Then, ∼1 × 10^5^ cells of each strain in 3 μL ddH_2_O were spotted on YPD + CuSO_4_ (1 mM) or YPD + phloxine B (5 μg/mL) medium and incubated at 30°C for four days.

### 3.7 Haploid and diploid cells of *C. glabrata* differ in fungal burdens in a mouse systemic infection model

Ploidy variations often regulate pathogenesis in fungal pathogens such as *C. albicans* and *C. auris* (11, 28). We next tested whether haploid and diploid cells of *C. glabrata* exhibited distinct levels of fungal burdens using a disseminated candidiasis mouse infection model. Generally, the order of fungal burden in different tissues was diploid > haploid, suggesting that the ploidy level is associated with pathogenesis or the ability of colonization in the host in *C. glabrata* (**Figure 6**). We did not include the hyperdiploid form in this assay since the genome of hyperdiploid cells are relatively unstable during infections.

**Figure 6.**
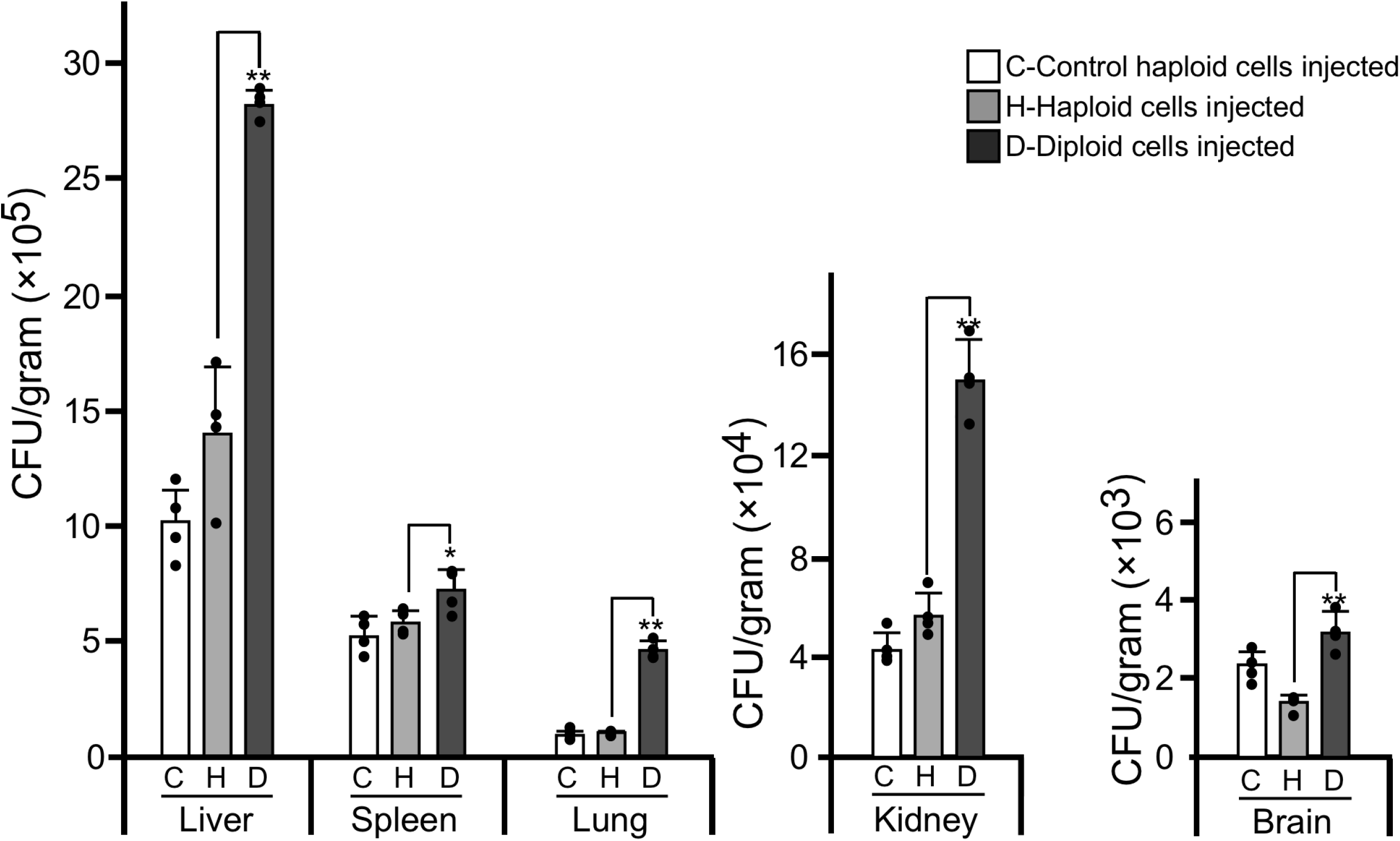
Fungal burdens of haploid and diploid *C. glabrata* cells in the mouse systemic infection model. Strain used: RJ155 (haploid control), FK83-1 (haploid), and FK83-2 (diploid). The haploid-only isolate (RJ155) served as the control. Four mice were used for each cell type. Each mouse was injected with 2 × 10^7^ cells of *C. glabrata* via the tail vein. After 24 hours of infection, mice were humanely killed, and fungal burdens of multiple organs were analyzed. The statistical significance of the differences between diploid and haploid values is indicated (*p<0.05, **p<0.01, Student’s *t*-test, two-tailed. *F*-test was used to compare and judge the homogeneity of variance in *t*-tests. The standardized effect size for *t*-tests was measured by Cohen’s *d*). Black dots shown in the figure represent the CFU data points of each mouse. Error bars represent standard deviations.

### 3.8 Haploid, diploid, and hyperdiploid cells of *C. glabrata* differ in susceptibility to the antifungal itraconazole

Itraconazole (ITC) is the first-line antifungal drug. We next tested the susceptibility to ITC in 16 clinical isolates with at least two ploidy forms. As shown in **Figure 7**, 56% of the samples (9/16) with different ploidy forms showed a significant difference in terms of MIC values. Generally, haploid cells had relatively higher MIC values, suggesting that ploidy variations could play a role in the regulation of antifungal resistance.

**Figure 7.**
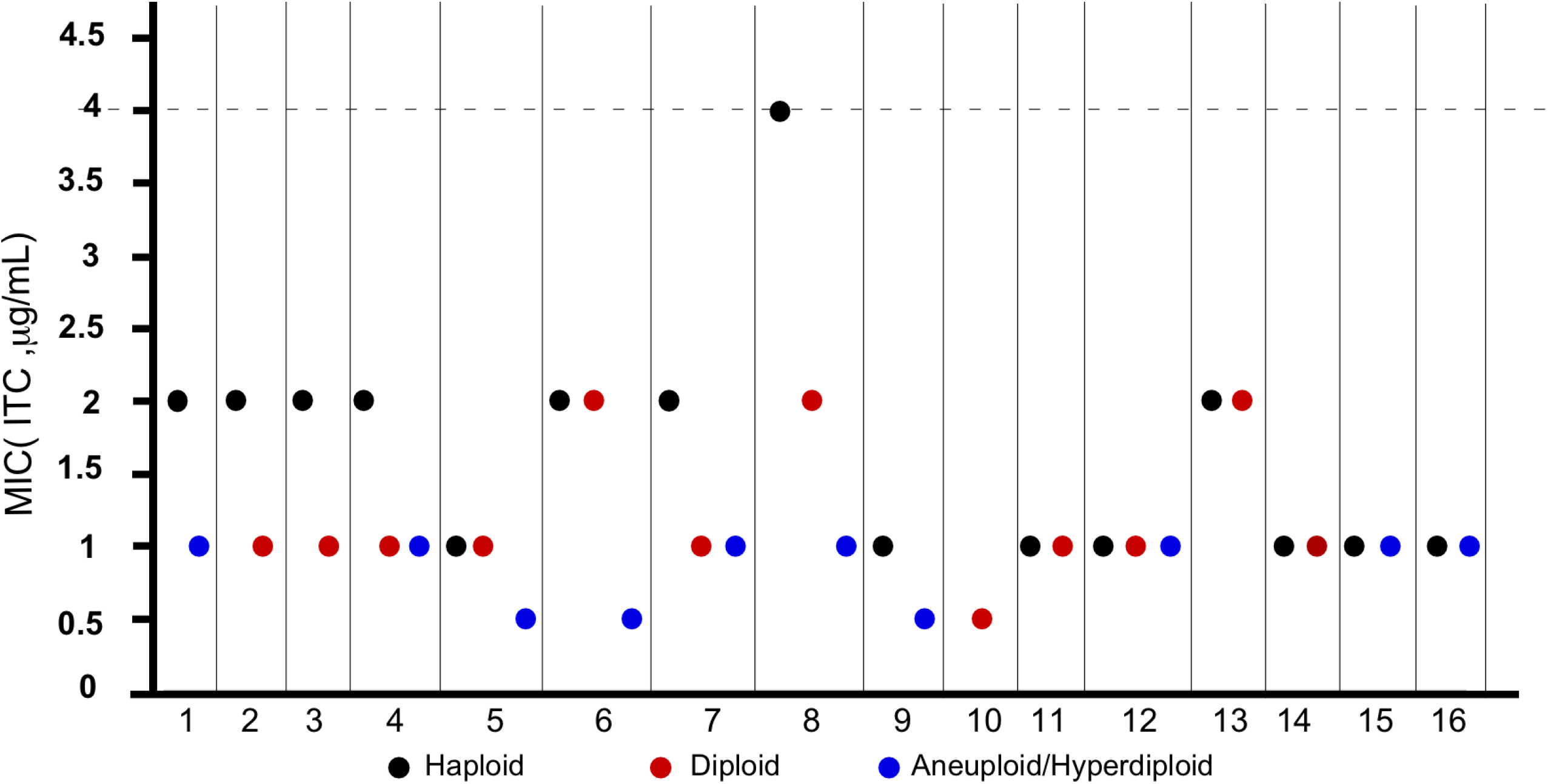
MICs of haploid, diploid, and hyperdiploid *C. glabrata* cells to itraconazole. MIC assays were performed according to NCCLS document M27-A2. A total of 38 strains derived from Renji Hospital (Shanghai), Shanghai Pulmonary Hospital, and the First Hospital of China Medical University (Shenyang) clinical isolates were tested. The strain information is presented in supplementary **Dataset S1**. Three biological replicates were assessed. The dotted line represents the epidemiological cut off value of itraconazole (ITC). *Candida parapsilosis* ATCC22019 and *Pichia kudriavzevii* ATCC6258 served as quality-control strains.

### 3.9 Haploid and diploid cells of *C. glabrata* differ in global transcriptional profiles

We next performed RNA-Seq analysis using strain FK83 to elucidate the potential mechanism underlying the biological differences between haploid and diploid cells of *C. glabrata*. We did not examine the global gene expression profile in aneuploid or hyperdiploid cells because their genomes were not stable. As shown in **Figure 8** and supplementary **Dataset S2**, a total of 750 genes were found to be differentially expressed between the two cell types (two-fold change cutoff, three replicates). Of them, 481 genes were upregulated in diploid cells, and 269 genes were upregulated in haploid cells. These differentially expressed genes are involved in many biological processes, such as DNA/RNA metabolism, chromatin and cyclin regulation, subcellular location, metabolism, and cellular adhesion. Genes associated with chromosome/nucleotide metabolism and the cell cycle were highly expressed in diploid cells (e.g., *TDA9, MSH5, DBP9, RAT1*, and *RTT106* for DNA metabolism and *NOP58, MTR4, RPO31, RPA190*, and *DBP2* for RNA metabolism). This gene expression profile suggested that diploid cells were more active in general metabolism than haploid cells, which was consistent with the relatively higher virulence of diploid cells during infection. Another possibility is that this gene expression difference could be due to the different growth phases of haploid and diploid cells. We observed a similar effect in haploid and diploid cells of *C. auris* (11).

**Figure 8.**
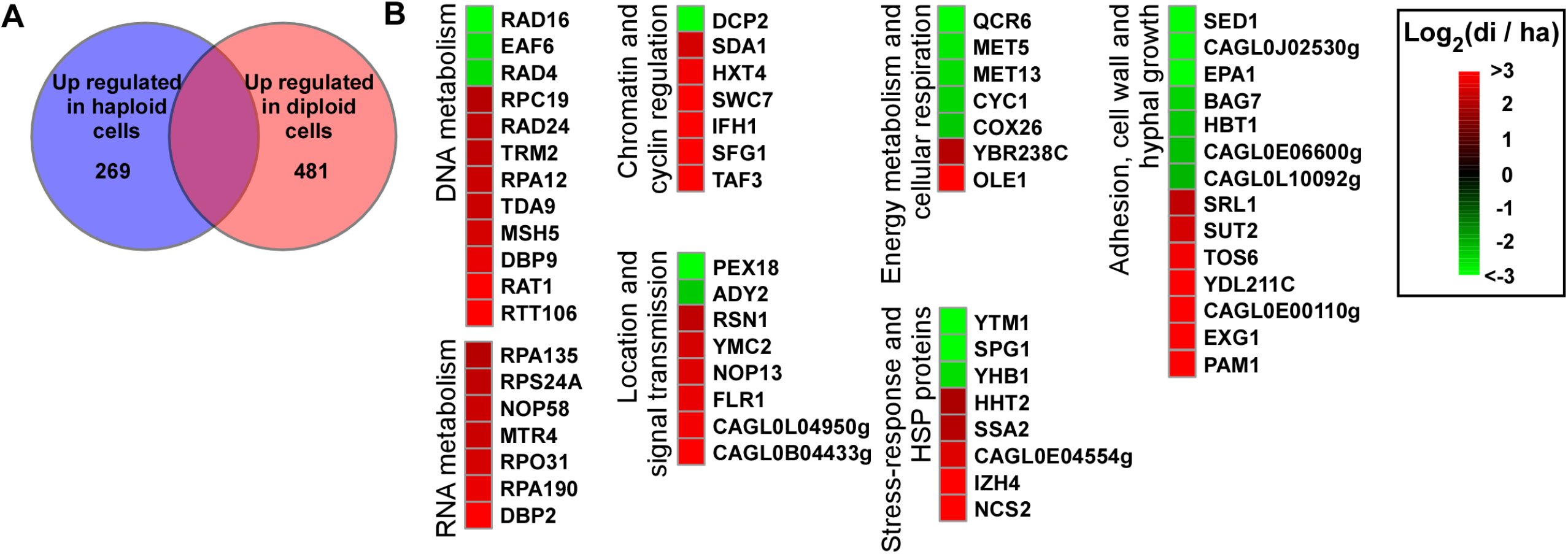
Different gene expression profiles in haploid and diploid cells of *C. glabrata*. RNA-Seq assays were performed using haploid and diploid cells of strain FK83. (A) Venn diagram of differentially expressed genes. A 2-fold difference cut off was used to define differentially expressed genes. (B) Representative genes that were differentially expressed. An R package heatmap was used to depict selected differentially expressed genes. Log_2_ (di/ha), Log_2_ (read counts of diploid cells/read counts of haploid cells). *C. glabrata* gene names or IDs align with the CGD database (http://www.candidagenome.org/).

Genes involved in energy metabolism and cellular respiration (e.g., *QCR6, MET5, MET13, CYC1*, and *COX26*) were upregulated in haploid cells. Moreover, some genes associated with the stress response, adhesion, cell wall and morphological transitions were also differentially expressed, suggesting that haploid and diploid cells differ in many biological aspects. For example, the haploid-enriched gene *EPA1* encodes an important adhesin that is required for adherence to host epithelial cells, biofilm formation, and antifungal resistance (29). Our findings indicate that haploid and diploid cells of *C. glabrata* exhibit unique global transcriptional profiles, which could contribute to their distinct responses to host and environmental conditions.

## 4. Discussion

Genomic plasticity is a rapid adaptive mechanism of human pathogenic fungi to the hostile environment. For example, aneuploidy and polyploidy are quite common in *C. albicans* and *Cryptococcus neoformans* (8). The model organism *S. cerevisiae* is able to undergo ploidy changes through both sexual reproduction and spontaneous shift under certain stressful conditions (10, 30). The switch between low and high ploidy forms promotes the generation of de novo mutations and diversity of phenotypes (9, 17). In this study, we report the discovery of ploidy variation and haploid-diploid switching in clinical isolates of *C. glabrata*. Haploid and diploid cells were also able to undergo spontaneous switching under both *in vitro* and *in vivo* conditions (**Figures 3 and 4**). More importantly, we isolated diploid cells of *C. glabrata* from original clinical samples, implying that ploidy variation could occur during host infections. The ploidy variation could be a general feature of clinical isolates. MLST analysis revealed two major genetic clusters of clinical *C. glabrata* isolates. Most Chinese strains, especially the strains with an altered ploidy form (diploidy or aneuploidy/hyperdiploid) belonged to clade A, whereas most *C. glabrata* strains from the USA and Iran belonged to clade B. The factors responsible for this genetic variation between different countries remain to be investigated.

Haploid, diploid, and hyperdiploid cells of *C. glabrata* differed in several aspects, including phenotype, antifungal resistance, and global gene expression profiles. On YPD + CuSO_4_ medium, haploid cells formed darker colonies than diploid and hyperdiploid cells (**Figure 5**). These chromogenic differences could reflect different redox states and abilities to respond to stresses. Consistently, transcriptional profile analysis indicated that a lot of genes involved in stress response were differentially expressed in *C. glabrata* haploid and diploid cells. Some isolates with different ploidy levels also differed in susceptibility to the antifungal ITC, suggesting that this change or underlying genetic variations could benefit *C. glabrata* survival under hostile conditions. In the disseminated mouse infection model, diploid cells of *C. glabrata* exhibited a higher fungal burden than haploid cells (**Figure 6**). Similar results have been reported with *C. auris* and *C. albicans*, the diploid cells of which are also more virulent than the haploid cells (11, 28). Consistently, we did not observe diploid-to-haploid switching in the mouse infection model, perhaps due to the increased *in vivo* colonization ability of diploid cells of *C. glabrata*. These biological differences could benefit fungal cells with distinct ploidy forms to better adapt to different ecological niches during infection or colonization of the host.

Taken together, ploidy variation could not only promote the rapid adaptation of *C. glabrata* to the changing environment but also benefit the evolution of new traits in the long term. The genome of clinical isolates of *C. glabrata* is extremely unstable (13), which might be due to the frequent change in ploidy forms. The increase in ploidy would result in the generation of aneuploidy and novel mutations. The spontaneous mutation rate of diploid cells has been reported to be much higher than that of haploid cells of *S. cerevisiae*, [26]. Moreover, *C. glabrata* has long been considered an “asexual” fungus. Ploidy switching among the haploid and diploid forms could function as an alternative biological process of sexual or parasexual reproduction. Our findings concerning the diploid form and ploidy variation in *C. glabrata* would shed new lights on its biology and pathogenesis.

## Funding

This work was supported by the National Key Research and Development Program of China (2021YFC2300400), National Natural Science Foundation of China (82172290 and 82002123 to HD, 31930005 to GH), and Shanghai Municipal Science and Technology Major Project (HS2021SHZX001).

## Conflicts of Interest

The authors declare no conflict of interest.

## Data availability statement

The RNA-Seq dataset has been deposited into the NCBI Gene Expression Omnibus (GEO) portal (accession # GSE173617).

## Supporting information

**Figure S1. Colony and cellular morphologies of two clinical samples (**RJ1271 and RJ1372 isolated from throat swabs) **of *C. glabrata* containing both haploid and diploid cells**. Four colonies of each sample showing distinct morphologies on CHROMagar medium were replated on YPD medium (containing 5 μg/mL of phloxine B, a red dye). The plates were then incubated at 30°C for four days. FACS assays were performed to determine the genomic DNA content. Scale bar for cells, 10 μm.

**Figure S2. The population structure of 194 *C. glabrata* isolates, including 70 isolates from four Chinese hospitals (analyzed in Figure 2) and 124 isolates from other countries** (Iran, USA, Japan, South American, and Europe). Multi-locus sequence typing (MLST) analysis was performed based on six genes (*FKS, LEU2, NMT1, TRP1, UGP1*, and *URA3*). The phylogenetic tree was constructed using the maximum-likelihood (ML) method. The General Time Reversible (GTR) model, Gamma distribution, and 1000 bootstraps were adopted. The isolates from Chinese hospitals are highlighted in red. The DNA sequence information for the 124 isolates from Iran, USA, Japan, South American, and Europe was downloaded from the NCBI database (accession number: KX187005-KX187304, AY771006-AY771209, KT763084-KT763323).

**Dataset S1. Clinical information of *C. glabrata* isolates**.

**Dataset S2. RNA-Seq dataset**.

